# The majority of Germany’s small agricultural streams are in poor ecological status

**DOI:** 10.1101/2023.10.27.564349

**Authors:** Julia von Gönner, Jonas Gröning, Volker Grescho, Lilian Neuer, Veit G. Hänsch, Benjamin Laue, Eva Molsberger-Lange, Elke Wilharm, Matthias Liess, Aletta Bonn

**Affiliations:** Helmholtz Centre for Environmental Research (UFZ), Department Ecosystem Services, Permoserstr. 15, 04318 Leipzig, Germany; Friedrich Schiller University Jena, Institute of Biodiversity, Dornburgerstr.159, 07743 Jena, Germany; German Centre for Integrative Biodiversity Research (iDiv) Halle-Jena-Leipzig, Puschstr. 4, 04103 Leipzig, Germany; Helmholtz Centre for Environmental Research (UFZ), Department System-Ecotoxicology, Permoserstr. 15, 04318 Leipzig, Germany; RPTU Kaiserslautern-Landau, Institute of Environmental Sciences, Fortstr. 7, 76829 Landau, Germany; Friends of the Earth Germany e.V. (BUND), Kaiserin-Augusta-Allee 5, 10553 Berlin, Germany; Saaletreff Jena, Beutnitzer Straße 5, 07749 Jena; Adolf-Reichwein-Schule, Heinrich-von-Kleist-Straße 14, 65549 Limburg an der Lahn, Germany; Ostfalia University of Applied Sciences, Am Exer, 38302 Wolfenbüttel, Germany; RWTH Aachen University, Institute of Ecology & Computational Life Science, Templergraben 55, 52056 Aachen, Germany

**Keywords:** small agricultural streams, European Water Framework Directive monitoring, pesticide pressure, citizen science, benthic invertebrate community composition, hydromorphology

## Abstract

Agricultural pesticides, nutrients, and habitat degradation are major causes of insect declines in lowland streams. To effectively conserve and restore stream habitats, standardized stream monitoring data and societal support for freshwater protection are needed. Here, we sampled 137 small stream sites across Germany, 83% of which were located in agricultural catchments, with more than 900 citizen scientists in 96 monitoring groups. Sampling was carried out according to Water Framework Directive standards as part of the citizen science freshwater monitoring program FLOW in spring and summer 2021, 2022 and 2023. The biological indicator SPEAR_pesticides_ was used to assess pesticide exposure and effects based on benthic invertebrate community composition. Overall, 58% of the monitored agricultural stream sites did not achieve a good ecological status in terms of macroinvertebrate community composition and indicated high pesticide exposure (SPEAR_pesticides_ status class: 29% ‘moderate’, 19% ‘poor’, 11% ‘bad’). The indicated pesticide pressure in streams was related to the proportion of agricultural land in the catchment area (R^2^=0.23, p<0.001). With regards to hydromorphology, monitoring results revealed that 65% of the agricultural stream sites failed to reach a good status. The data base produced by citizen science groups was characterized by a high degree of accuracy. This was quantified by a comparative survey of SPEAR_pesticides_ index (R^2^=0.79, p<0.001) and hydromorphology index values (R^2^=0.72, p<0.001) by citizen scientists and professionals. Such citizen-driven monitoring of the status of watercourses could play a crucial role in monitoring and implementing the objective of the European Water Framework Directive, thus contributing to restoring and protecting freshwater ecosystems.

**Graphical abstract:** 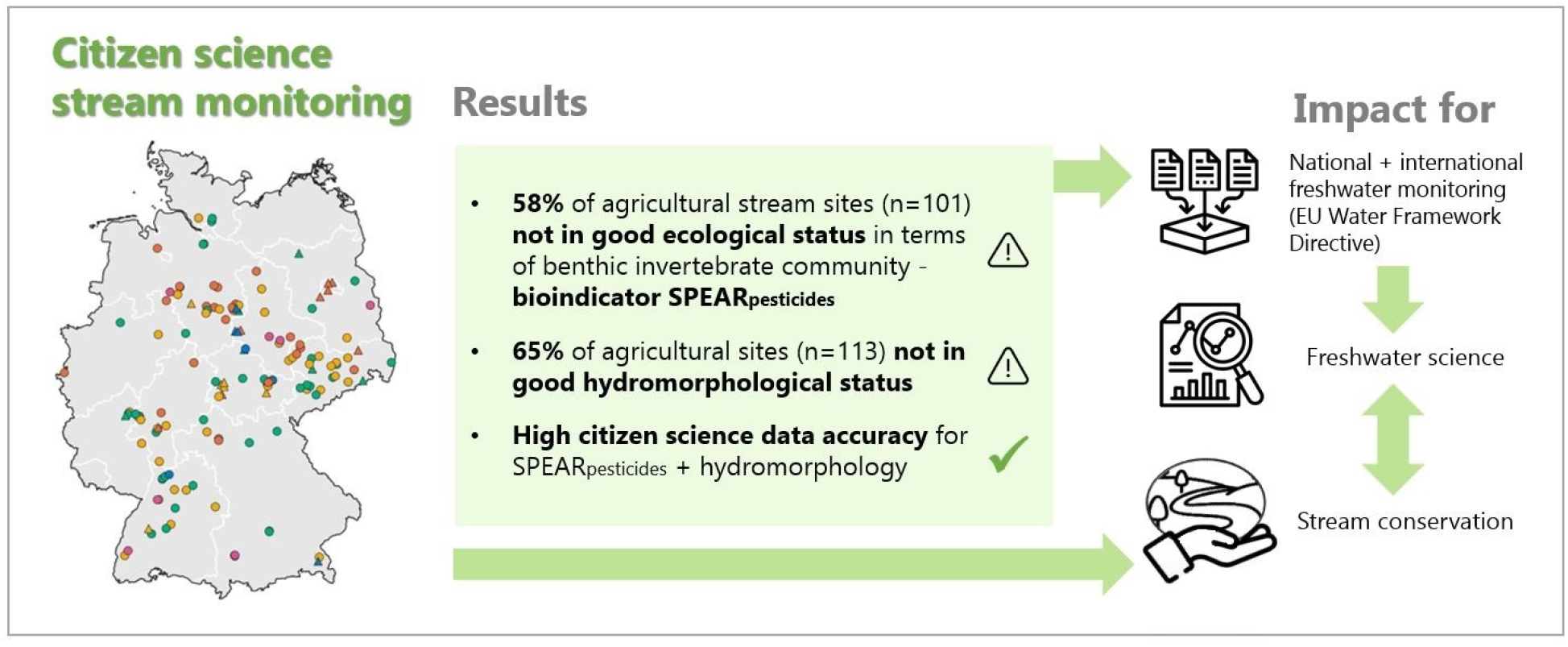

**Highlights:**

- Assessment of ecological status of small agricultural streams in Germany
- Pesticides affected invertebrates (bioindicator SPEAR) in 58% of agricultural streams
- Failure to reach good hydromorphological status in 65% of agricultural streams
- Citizen science monitoring achieves high data accuracy
- Citizen science can support European Water Framework Directive monitoring

## 1. Introduction

### 1.1 Current status of river protection and monitoring in Europe

With the Water Framework Directive (WFD, European Commission, 2000) and the European Green Deal (European Commission, 2019a), the European Union has adopted ambitious policies to protect freshwater ecosystems. The WFD’s goal is to conserve or restore the good ecological status of all surface waters and to restore a good ecological potential in heavily changed water bodies (EPA, 2006). A ‘good ecological status’ according to WFD is achieved in rivers and streams when hydromorphology, physico-chemical status and biotic communities (i.e. macrophytes/algae, macroinvertebrates and fish) are in a near-natural condition specific to the relevant ecoregion and river type. Several ecosystem services essential for human well-being, such as water storage and purification, conservation of (semi)-aquatic biodiversity and recreational qualities, are closely linked to good status of rivers and streams (Schäfer et al., 2012; Chung et al., 2021).

To date, however, member states have repeatedly failed to meet freshwater protection targets (EEA, 2018; European Commission, 2019b; IPBES, 2019). As evidenced by recent freshwater monitoring studies and agency reports, about 60% of river sample sites in Europe (EEA, 2018) and more than 80% of rivers sampled in Germany are in a poor ecological status (Liess et al., 2021; UBA, 2022). Water Framework Directive monitoring is generally carried out in rivers and streams with a focus on catchment areas >100km^2^. Pesticide and nutrient inputs, as well as human-made hydromorphological and climatic changes are the main drivers of the poor ecological status and observed declines in vulnerable insect diversity in streams (Vörösmarty et al., 2010; Liess et al., 2021; Wolfram et al., 2021). Due to these anthropogenic stressors, aquatic biodiversity (Beketov et al., 2013) and stream-based ecosystem services such as leaf and organic matter decomposition (Schäfer et al., 2012; Böck et al., 2018) are strongly impaired.

Small streams account for two thirds of the entire river network in Germany (Meyer et al., 2007; BfN, 2021) and thus play an important role for freshwater and biodiversity conservation. Due to their small water volume and in many cases, proximity to agricultural land use, they can be especially affected by pesticide and nutrient inputs (Halbach et al., 2021; Szöcs et al., 2017). There is a lack of systematic large-scale monitoring data on small streams, however, since these small streams with catchment areas below 30km^2^ are only rarely monitored and streams below 10km^2^ are not taken into account in the official WFD monitoring scheme (Wick et al., 2019; Weisner et al., 2022).

In order to reduce the drastic loss of biodiversity, habitat and ecosystem services at EU level, and to ensure the implementation of the EU Green Deal, the European Commission has recently adopted proposals for the ‘Nature Restoration Law’ (NRL, European Commission 2022a) and ‘Regulation on the sustainable use of plant-protection products’ (SUR, European Commission, 2022b). These state that at least 30% of degraded habitats in terrestrial, coastal, freshwater and marine ecosystems should be restored to a good ecological status by 2030. The use of pesticides and the associated ecological and health risks are to be reduced by 50% by 2030, and pesticides are to be completely banned in sensitive areas, although scientists fear that effective implementation of the planned measures is at risk (Pe’er et al., 2023). In order to achieve an evidence-based and accountable implementation of the Water Framework Directive, the Restoration Law and the Sustainable Use Regulation, the ecological status of freshwater ecosystems of all sizes needs to be assessed on a broad spatio-temporal scale. The data can then be used to identify effective restoration priorities and measures.

### 1.2 Benthic invertebrates as biological indicators for pesticide exposure

Benthic invertebrates are widely used as biological indicators to assess the ecological status of rivers and streams (Chessman et al., 2007; Friberg et al., 2011). They are sensitive to several ecological stressors, relatively easy to sample, and have been shown to be suitable biological indicators in the context of citizen science stream monitoring projects (Storey et al., 2016; Brooks et al., 2019; Moolna et al., 2020).

The biological indicator SPEAR_pesticides_ has been developed to quantify ecological effects of pesticide exposure in agriculturally influenced streams based on macroinvertebrate community composition (Liess & v.d. Ohe, 2005; Liess et al., 2021). SPEAR is a trait-based indicator based on the relative abundance of pesticide-sensitive macroinvertebrate taxa at a stream site (Liess & v.d. Ohe, 2005). Based on four ecological functional traits (physiological sensitivity to pesticides, generation time, life cycle or hatching time, and ability to migrate and recolonize), each macroinvertebrate taxon is categorized as either ‘SPEcies At Risk’ (SPEAR) or ‘SPEcies not At Risk’ (SPEnotAR). The SPEAR indicator has been shown to be a suitable method for identifying pesticide exposure and establishing dose-effect relationships at large spatial scales. It mainly reacts to pesticide exposure and is mostly independent of other stressors such as oxygen deficiency or nutrient load (Liess et al., 2008; Knillmann et al., 2018; Liess et al., 2021). SPEAR_pesticides_ is used for pesticide indication in the German WFD stream assessment (LAWA and UBA, 2022). Previous research has shown that the indicator provides accurate results with macroinvertebrate data whose taxonomic resolution is limited to family level (Beketov et al., 2009; Liebmann et al., 2022) and is therefore well suited for citizen science stream monitoring (von Gönner et al., 2023a).

### 1.3 Potentials of citizen science

Effective freshwater monitoring and protection is a multi-faceted, challenging task. It requires not only scientific expertise and practical knowledge, but also the involvement of local communities, different stakeholders and the compliance of large parts of society with freshwater protection measures (EPA, 2006; Carvalho et al., 2019; BMUV, 2023).

Citizen science, the active participation of interested citizens in research processes, holds great potential to advance ecological stream monitoring and restoration (Storey et al., 2016; Brooks et al., 2019; von Gönner et al., 2023a). By collecting data on riverine species or the status of freshwater ecosystems, citizen scientists can provide new knowledge as a basis for scientific analyses (Bowler et al., 2021; Maasri et al., 2022). Simultaneously, research has shown that citizen science projects in water monitoring can raise awareness for freshwater ecosystems and their services (Storey et al. 2016). Citizen science can foster social license for conservation (Kelly et al., 2019), promote stewardship for local streams and enable citizens to engage in stream protection (Brooks et al., 2019; Huddart et al., 2016; Edwards et al., 2018).

In Germany, freshwater ecology is also anchored in the upper secondary school curriculum, but rarely practiced in the field. Therefore, citizen science could be a valuable tool for raising students’ awareness of aquatic ecosystems and their stressors, and for increasing their self-efficacy in freshwater conservation through research-based, hands-on outdoor learning (see Ballard et al., 2017).

### 1.4 Current study

To co-create new knowledge on the ecological status of streams, we developed the citizen science stream monitoring program FLOW in Germany (https://flow-projekt.de; von Gönner et al. 2023a). The FLOW program provides learning material, training, support, field equipment and a digital data management system for citizen scientists to generate WFD compliant data on stream hydromorphology and macroinvertebrate community composition. FLOW specifically focuses on small streams with catchment sizes below 30 km^2^ to fill data gaps and complement the official WFD stream monitoring. To assess pesticide exposure at the stream sample sites, the macroinvertebrate data is used to calculate the bioindicator SPEAR_pesticides_. FLOW mobilized a total of 96 citizen science groups between 2021 and 2023 with over 900 participants who assessed the ecological status of their local stream sites. Our previous study on the accuracy of citizen science data (von Gönner et al., 2023a) showed on a smaller sample of 28 streams that personally trained citizen scientists can correctly identify macroinvertebrates to the family level and provide valid data on the SPEAR_pesticides_ index and stream hydromorphology. Here, we again provide detailed data on the accuracy of citizen science freshwater sampling data compared to professional assessments of SPEAR_pesticides_ and hydromorphology index values before analyzing the data.

In this study, based on three years of citizen science stream monitoring in Germany,

- we assess the ecological status of small streams in Germany and the proportion of stream sample sites that achieves a good ecological status with regards to benthic invertebrate community, hydromorphology and physico-chemical status
- we also assess the impact of land cover in the analyzed stream catchments, and how agricultural land cover is related to pesticide pressure indicated by SPEAR_pesticides_ index values.

Our study provides the largest citizen science study of small agricultural stream sites in Germany. Based on the results, we discuss possible reasons for stream sites not achieving good ecological status. We provide an outlook on how citizen science freshwater monitoring can contribute to greater accountability for agreed government conservation targets and, potentially, support restoration efforts.

## 2. Methods

### 2.1 Study design and site selection

We used a two-tiered approach for selecting stream sample sites. For the first FLOW monitoring campaign in 2021 (pilot phase), we selected 30 agricultural stream sites in central Germany with catchment sizes up to 30 km^2^ to test and fine-tune the citizen science monitoring alongside professional monitoring (for details see von Gönner et al., 2023a). In the 2022 and 2023 monitoring campaigns, we extended the FLOW monitoring across Germany (see Fig. 2). Stream sites were chosen according to the following criteria: catchments smaller than 30 km^2^; priority to agricultural streams with at least 20% agricultural land cover in their catchments; possibility to assess streams with less than 20% agricultural land cover in catchment as a comparison baseline (referred to as non-agricultural streams); as few urban areas as possible in catchment and no wastewater treatment plants upstream to focus on agricultural diffuse pollution (Liess et al., 2021). Catchments of the analyzed stream sites were characterized by a gradient of agricultural land (mean agricultural land cover 53% ± 29%, for detailed site characteristics see SI Fig.1; SI Tab.1).

### 2.2 Citizen science training

Over the years, we recruited 96 citizen scientist groups (42 local environmental NGO groups, 26 local citizen initiatives, 18 senior high school classes, and 10 angling clubs), with a total of over 900 participants. A large majority of the participating citizen scientists were interested newcomers with little to no prior experience in ecological stream assessment. To foster high quality monitoring, several local freshwater experts (with in-depth taxonomic or ecological knowledge gained through long-term voluntary engagement) participated in the citizen science monitoring as group leaders. The term ‘professionals’ is used in this study to refer to experienced ecologists who acquired expertise in limnology as full-time researchers.

In the pilot phase of the FLOW project (2021, von Gönner et al., 2023a), all citizen science groups were trained directly by the FLOW team and also accompanied during the stream monitoring events to clarify open questions and help with any problems. In 2022 and 2023, we used a multiplier approach (‘Train-the trainer’) to ensure that all citizen science groups across Germany were well-prepared for the stream assessments: First, all citizen scientists participated in a 2-hour online training where we introduced and discussed the stream assessment methods. Subsequently, all citizen science group leaders attended an additional full day on-site training on macroinvertebrate identification down to family level. During the identification training, citizen science group leaders learned about the distinguishing characteristics of the major macroinvertebrate families and practiced sorting and identifying voucher specimens using a stereo microscope. For review and further practice, we provided additional learning materials for all participants: a project booklet with field protocols and background information on stream ecology, six short video tutorials on the FLOW stream monitoring methods, an identification booklet characterizing about 200 native macroinvertebrate taxa with photos and illustrations, and an online quiz on macroinvertebrate identification and hydromorphological assessment (for details on the training materials, see von Gönner et al. 2023a and SI Tab.2). After the identification training, citizen science group leaders organized preparatory meetings with their local groups to practice invertebrate identification. Some group leaders contacted experienced biologists in their region for assistance with the field work. This proved to be a very effective method of supporting citizen scientists in the field with sampling and identification.

### 2.3 Data collection and preparation

Citizen scientists assessed benthic invertebrate community composition, hydromorphology, and physico-chemical status at a total of 137 stream sites in 2021, 2022 and 2023, including 113 agricultural sites and 24 non-agricultural sites. Most sites (57%) were sampled once in one of the three monitoring years by a group of 5 to 15 trained citizen scientists between April and early July, the main pesticide application period for most crops (Szöcs et al. 2017). Citizen science groups with sufficient time capacity sampled their sample sites twice (in April and June of one year, 23% of sample sites) or more frequently (in two or three subsequent years, 20% of sample sites) to document seasonal changes and changes over time.

For the analysis in this study, we selected the most recent sampling result from the sampling period of May and June for all sites that were sampled two or more times. Previous analyses (Liess & v.d. Ohe, 2005) had shown that macroinvertebrate community composition in early summer best reflected current pesticide exposure. Including mean values for sites with multiple sampling events did not significantly change the result patterns. For the analysis of SPEAR_pesticides_ bioindicator values, we excluded all sites where macroinvertebrate communities were severely affected by a lack of flow (dried out or flow velocity < 0.05 m/sec) in the period from April to July so that accurate bioindication was not possible (Liess et al. 2021). This resulted in a sample of n = 120 sites for the analysis of SPEAR_pesticides_ values (i.e., 101 agricultural and 19 non-agricultural sites).

Citizen scientists sampled benthic invertebrates using standardized multi-habitat sampling according to the WFD: they first recorded stream bed substrates (Meier et al., 2006) and documented the distribution of substrate types on a percentage basis (smallest unit 5%). A total of 20 subsamples was then divided proportionally between the occurring substrate types present: Each subsample substrate unit (5%) was sampled by kick sampling ten times using a net with a surface of 0.0625m^2^ and a mesh size of 0.5 mm (Liess et al., 2021). Sampled invertebrates were separated from the coarse organic debris using a kitchen sieve or, where available, a column sieve set. Tweezers were used to sort the invertebrates into white trays. Citizen scientists then identified and counted the sampled invertebrates at least to family level using stereo microscopes with 20-fold magnification. The identification of live animals was conducted on-site directly at the stream site. Thus, citizen scientists had only one afternoon to complete the invertebrate sorting, counting, and identification. A subsample of n = 81 macroinvertebrate samples were preserved in 90% ethanol and determined to the highest possible taxonomic resolution in the laboratory by professional freshwater scientists. To assess stream hydromorphology and derive corresponding index values, citizen scientists used an illustrated and annotated version of the official protocol for small to medium-sized streams of the German Water Working Group of the Federal States (LAWA, 2019, see von Gönner et al., 2023a). All hydromorphological criteria required by the WFD were quantified, including meandering of the watercourse, variation in stream depth and width, flow diversity as well as bed habitat structure, riparian conditions and land use within a 100m stretch of the stream sampling section (European Commission, 2000).

For additional information on stream stressors, citizen scientists also measured physico-chemical water parameters (i.e. nitrite, nitrate, phosphate, pH, water temperature, dissolved oxygen, electrical conductivity, flow velocity) once per site in the afternoon of the citizen science sampling day. Repeated citizen science measurements were not possible in most cases for logistical reasons (see SI Tab.3 for information on measuring devices).

In 2021, citizen scientists entered their monitoring data into prepared Excel spreadsheets or the SPEAR calculator (https://www.systemecology.de/indicate/) to generate the relevant index values and to determine biological, morphological and physico-chemical status classes according to the WFD (von Gönner et al., 2023a). In 2022 and 2023, citizen scientists used the FLOW web application hosted by the Helmholtz Centre for Environmental Research (https://webapp.ufz.de/flow/) to upload and evaluate their stream monitoring data. This digital citizen science data management system is used to generate SPEAR and hydromorphology index values and to assign the monitoring results for all three quality elements to one of the five water status classes according to the WFD. It also serves as a platform for data visualization, download and archiving. For the present study, all SPEAR_pesticides_ values were recalculated according to the online SPEAR calculator version 2023.08.

### 2.4 Assessment of citizen science data accuracy

To assess citizen science invertebrate identification accuracy, we re-identified a subsample of n = 81 citizen science invertebrate samples (n = 30 samples from 2021, n = 30 samples from 2022 and n = 21 samples from 2023) in the laboratory to generate corrected invertebrate taxa lists. To verify citizen science hydromorphology assessments, we re-assessed the six subcategories of hydromorphology (i.e. water course, longitudinal profile, transverse profile, bed structure, bank structure, land use) for the same stream sites (n = 79) based on voucher photos taken by the citizen scientists. We then generated professional hydromorphology index values to compare them to the citizen science hydromorphology data. For measurements of physico-chemical water parameters no professional reference data was available.

We calculated linear regressions for SPEAR_pesticides_ and hydromorphology index values generated by the citizen scientists and the professionals, checking residuals for normality and homoscedasticity. To quantify bias, we additionally calculated the Concordance Correlation Coefficient (CCC) for SPEAR_pesticides_ and hydromorphology index values provided by citizen scientists and professionals using the epi.ccc function from the epiR R package (version 2.0.62, Stevenson and Sergeant, 2023).

The re-identification of n = 81 citizen science macroinvertebrate samples in the laboratory showed that the rate of correct citizen science identifications was high at the family level (Mean = 84%, SD = 13.4%, Fig. 1A). For the re-identified citizen science invertebrate samples, we found that citizen science and professional SPEAR_pesticides_ values were highly correlated (R^2^=0.79, p < 0.001, CCC = 0.87, Fig. 1A). 64% of the re-examined stream sites were rated with the same SPEAR status class by both citizen scientists and professionals, while 33% of the sites were rated one SPEAR class apart. On average, citizen scientists rated SPEAR index slightly more positively (mean index value = 0.55, SD = 0.31) than professionals (mean index value = 0.50, SD = 0.26, W = 2117, p <0.05, see also SI Fig. 4). Overall, citizen science and professional SPEAR assessments agreed in 84% of the cases on whether a stream achieved a good status in terms of pesticide exposure (i.e. classification as SPEAR status class I or II).

**Fig. 1.**
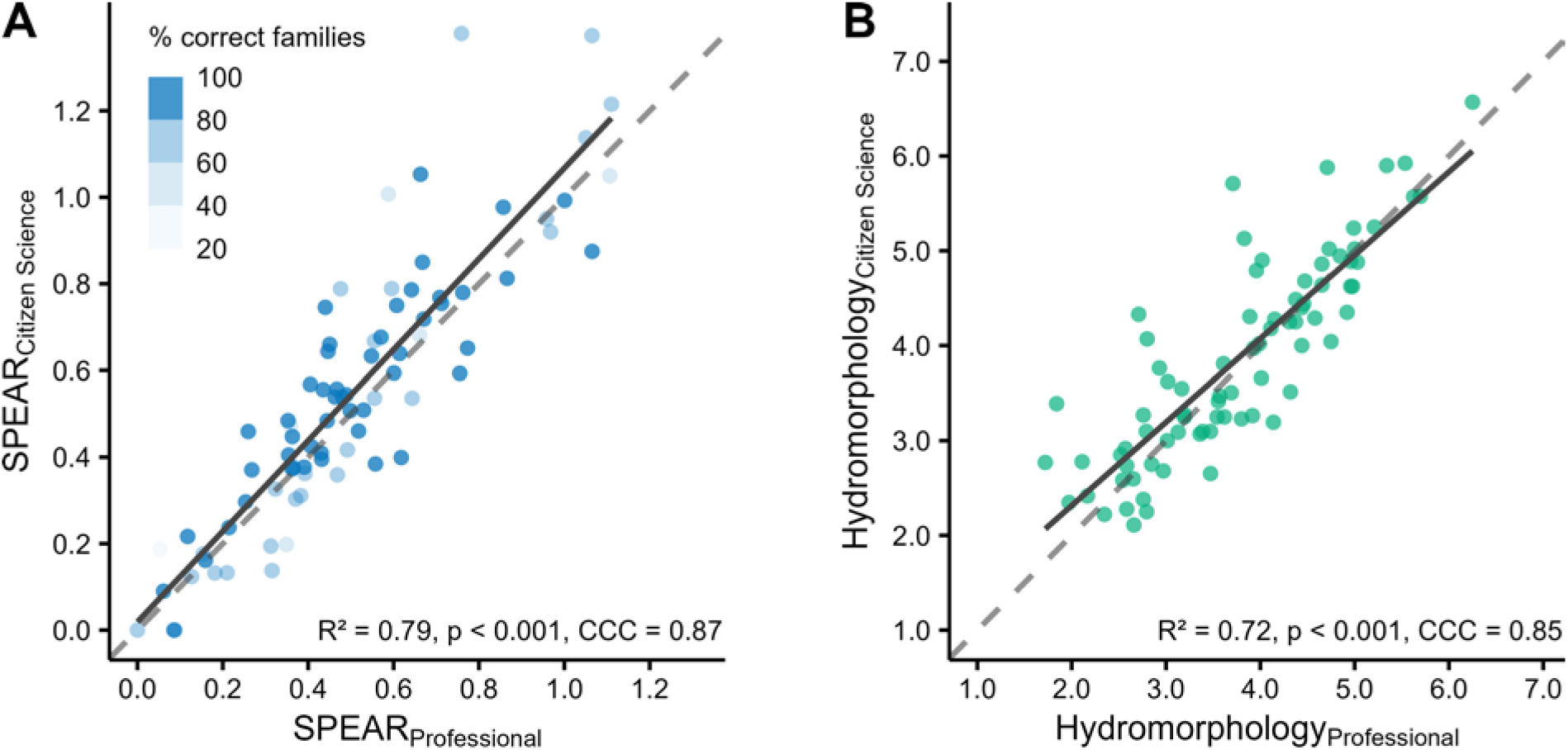
Correlation between citizen science and professional stream assessments for A. SPEARpesticides index values (n = 81) and B. hydromorphology index values (n = 79). SPEARpesticides index values are colored according to the percentage of correct citizen science identifications of invertebrate families per stream site. The solid line represents the regression line, the dashed line the 1:1 line. CCC = Concordance Correlation Coefficient.

**Fig. 2:**
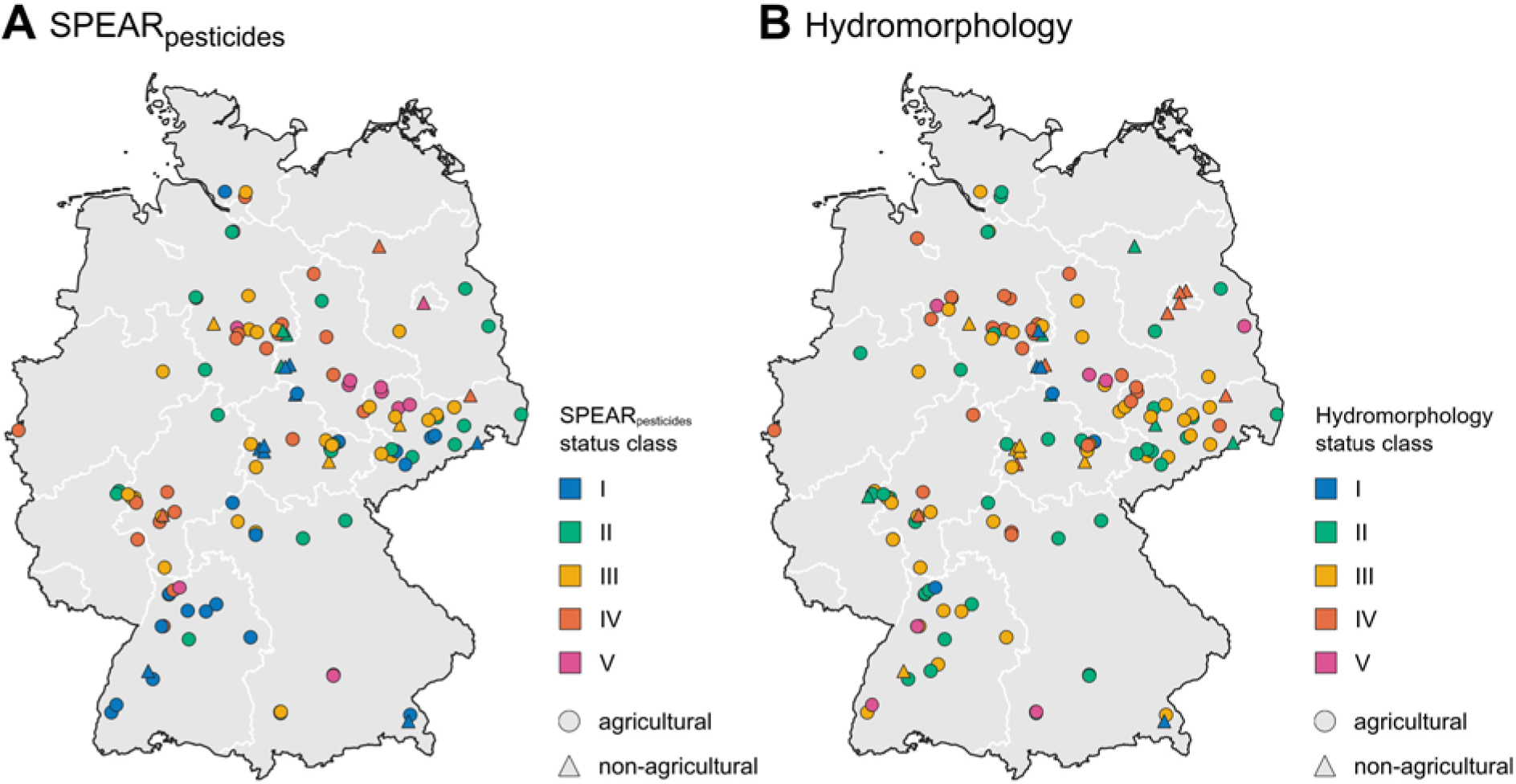
Map of Germany showing citizen science A. SPEAR_pesticides_ (n = 120) and B. hydromorphology status class (n = 137). For the analysis of SPEAR_pesticides_, we excluded n = 17 sites with very low flow velocity (<0.05m/sec) where accurate bioindication was not possible (see section 2.3). The symbols are colored according to the ecological status class. Agricultural sites are shown as a dot, non-agricultural sites as a triangle.

The re-examination of citizen science hydromorphology data for 79 sites revealed that citizen science and professional hydromorphology index values were highly correlated (R^2^=0.72, p < 0.001, CCC = 0.85, Fig. 1B). In detail, we found that 65% of the stream sites were rated with the same status class by both citizen scientists and professionals, while 35% were rated one status class apart (SI Fig. 5). We found no significant difference between citizen science hydromorphology index values (Mean = 3.85, SD = 1.05) and professional index values (Mean = 3.77, SD = 1.02, W = 1741, p = 0.43). Overall, citizen science and professional hydromorphology assessments agreed in 85% on whether a stream achieved a good ecological status according to WFD.

### 2.5 Field data analysis

First, we performed a descriptive analysis of index values and status classes for SPEAR_pesticides_ and hydromorphology as well as physico-chemical parameters, separately for agricultural and non-agricultural sites. The classification of the sites was based on the proportion of agricultural land-use in their catchments. Sites with >20% agricultural area in their catchment were classified as ‘agricultural’, sites with <20% agricultural area as ‘non-agricultural’ (Liess et al., 2021). To determine agricultural land cover, the catchments were delineated from a digital elevation model (EEA, 2016) and intersected with land-use data (EEA, 2020). We performed linear regressions to assess the influence of arable land in the catchments of the stream sites on SPEAR_pesticides_. Residuals were checked for normality and homoscedasticity. Single measurements of physico-chemical parameters were compared with regulatory thresholds from the German Surface Waters Ordinance (BGBl, 2016) and LAWA (1998) depending on stream type. We tested for differences between measurements in agricultural and non-agricultural streams using t-test or Wilcoxon test for normally and non-normally distributed data, respectively.

All statistical analyses were performed using R software (version 4.3.1, R Core Team 2023). Graphical visualizations were created using the ggplot2 R package (version 3.4.2, Wickham, 2016). Maps were produced using QGIS (version 3.32.1, QGIS Development Team 2023).

## 3. Results

### 3.1 Macroinvertebrate communities and SPEARpesticides index

In total, we analyzed invertebrate communities and SPEAR_pesticides_ index values for 120 stream sites (Fig. 2). Of the 101 agricultural stream sites, 58% did not achieve good ecological status with respect to macroinvertebrate communities according to the SPEAR_pesticides_ bioindicator. At these sites, SPEAR_pesticides_ index values were assigned to status classes III ‘moderate’ (29%), IV ‘poor’ (19%) or V ‘bad’ (11%), indicating that macroinvertebrate communities were negatively affected by pesticide inputs (Fig. 3A). Of the 19 non-agricultural stream sites, 37% did not reach good ecological status and were assigned to SPEAR_pesticides_ status classes ‘moderate’ (16%), ‘poor’ (16%) or ‘bad’ (5%).

**Fig. 3:**
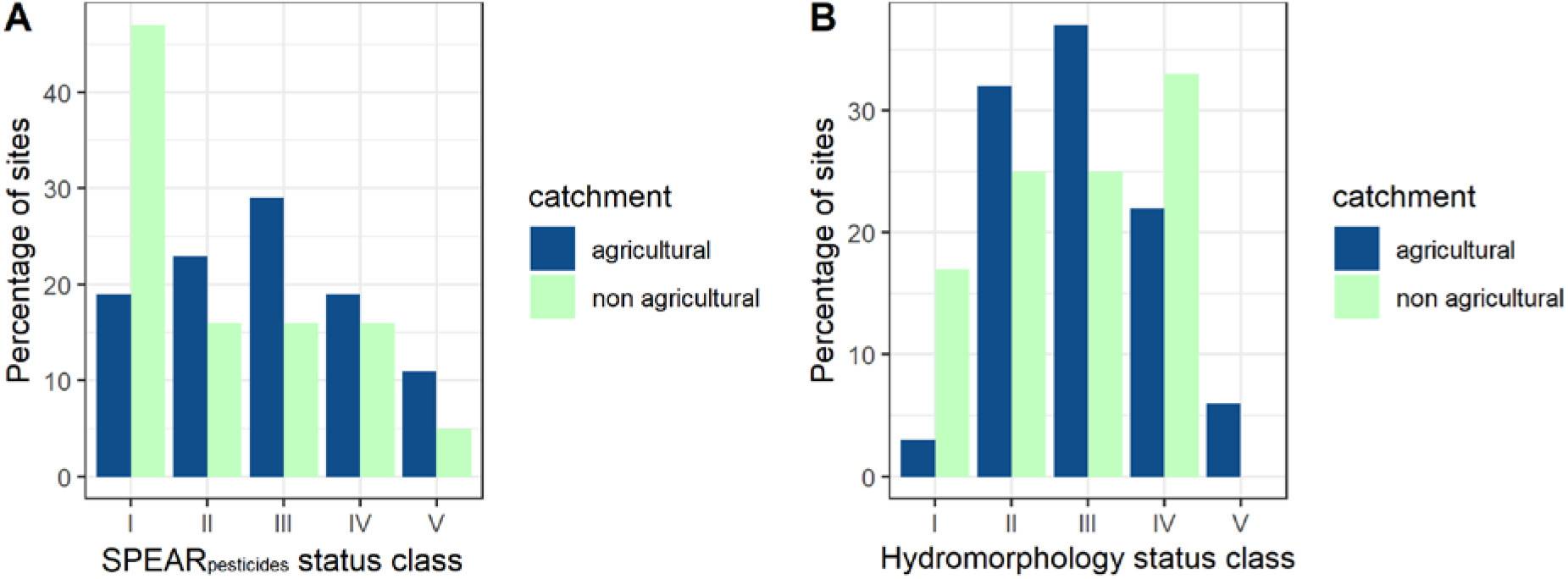
Proportion of A. SPEAR_pesticides_ status classes (n_total_ = 120; 101 agricultural catchments, 19 non-agricultural catchments) and B. hydromorphology status classes (n_total_ = 137; 113 agricultural catchments, 24 non-agricultural catchments). For the analysis of SPEAR_pesticides_, we excluded n = 17 sites with very low flow velocity (<0.05m/sec) where accurate bioindication was not possible (see section 2.3).

Catchment land cover mattered. SPEAR_pesticides_ showed a significant association with the proportion of arable land cover within the catchments (R^2^ = 0.23, p < 0.001). SPEAR_pesticides_ values decreased as the proportion of arable land increased, indicating higher pesticide pressures (Fig. 4).

**Fig. 4.**
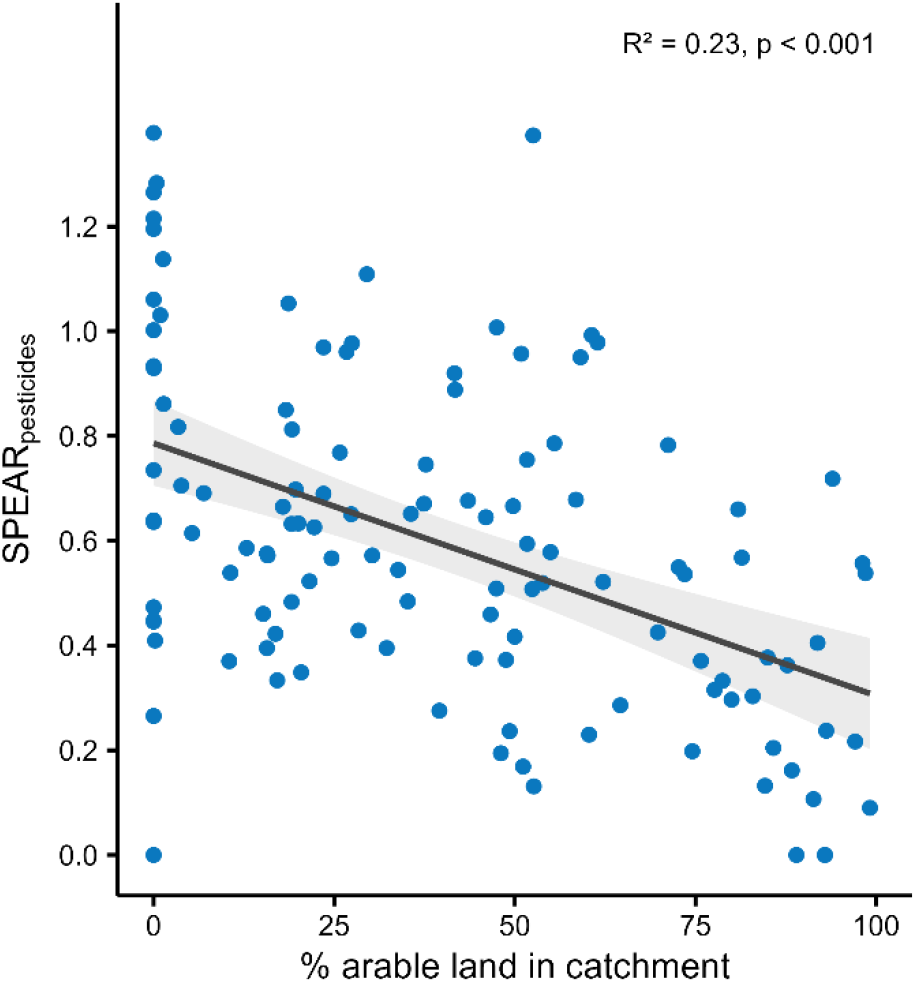
Relationship between the share of arable land and SPEAR_pesticides_ index values. A lower SPEAR_pesticides_ index indicates a higher pesticide pressure. The black line represents the regression line. The grey band corresponds to the 95% confidence interval.

### 3.2 Assessment of stream hydromorphology

For stream hydromorphology, 65% of the monitored agricultural stream sites (n = 113) did not reach a good ecological status (Fig. 3B), classified as status classes III ‘moderate’ (37%), IV ‘poor’ (22%) and V ‘bad’ (6%). Of the 24 non-agricultural sites, 58% did not achieve good ecological status and were assigned to hydromorphology status classes ‘moderate’ (25%) or ‘poor’ (33%). For details on the assessment of hydromorphology subcomponents, see SI Fig. 3.

### 3.3 Assessment of physico-chemical water status

For physico-chemical parameters, individual measurements exceeded at least one threshold for a good ecological status in 96% of the streams investigated (Fig. 5). Most frequently, the nutrients phosphate (79%), nitrate (73%), nitrite (42%) and ammonium (32%) concentrations exceeded the respective threshold values. Oxygen concentrations did not meet the criteria for a good ecological status in 25% of the streams. Less frequently, the regulatory limits for pH (7%) and chloride (3%) were exceeded.

**Fig. 5:**
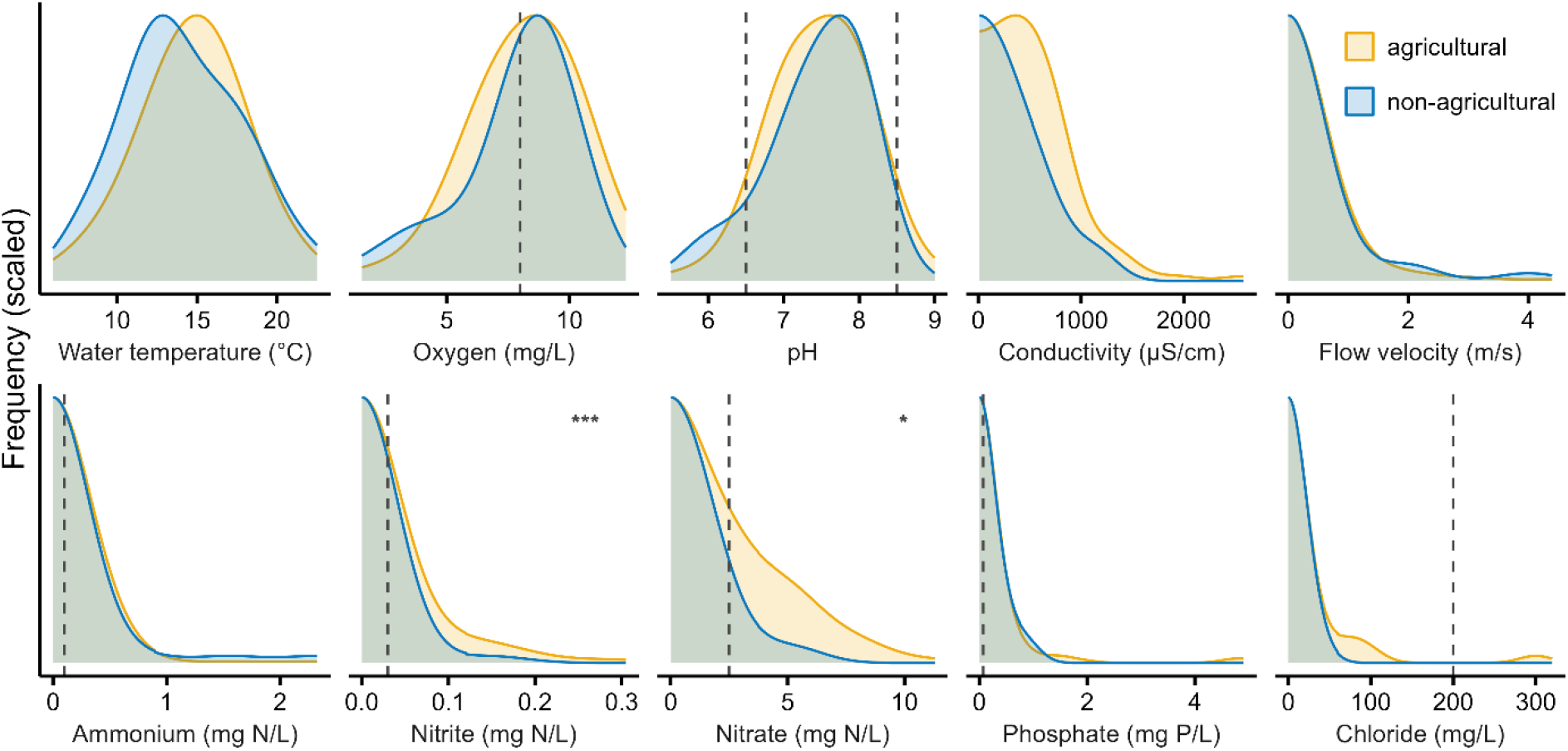
Distribution of citizen science measurements of physico-chemical parameters across the 137 sample sites. Asterisks indicate significant differences between measurements in agricultural and non-agricultural streams ^(*^ < 0.5, ^***^ < 0.001). Regulatory thresholds associated with good physico-chemical status (BGBl, 2016 or LAWA, 1988) are indicated with dashed lines. For visualization purposes, the threshold values are shown for the most common stream type in our investigations (type 5: coarse material-rich, siliceous low mountain stream). Thresholds for other stream types used to calculate exceedance frequencies may differ.

On average, nitrate concentration were two times higher in agricultural streams than in non-agricultural streams and nitrite concentrations even three times higher (Nitrate concentrations agricultural: mean = 2.76 mg N/L, non-agricultural: mean = 1.32 mgN/L, W = 1669, p < 0.05; Nitrite concentrations agricultural: mean = 0.041 mgN/L, non-agricultural: mean = 0.013 mgN/L, W = 1880, p < 0.001). Water temperatures (mean = 14.6 °C), oxygen concentrations (8.2 = mg/L), conductivity (523 μS/cm) and flow velocity (mean = 0.5 m/s) were similar in agricultural and non-agricultural streams. The nutrients ammonium (mean = 0.17 mgN/L) and phosphate (mean = 0.36 mgP/L) as well as chloride (25 mg/L) also showed no significant differences.

## 4. Discussion

Based on a large sample of lowland and highland stream sites in agricultural and non-agricultural catchments across Germany, our study provides the first comprehensive citizen science-based insight into the ecological status and pesticide exposure of small water bodies in Germany.

Analyzing citizen science stream monitoring data for a total of n = 137 stream sites across 13 German federal states, we found that 58% of the monitored agricultural sample sites did not meet good ecological status in terms of benthic invertebrate communities according to Water Framework Directive (WFD). At these stream sites, SPEAR_pesticides_ bioindicator values were classified as status classes III ‘moderate’, IV ‘poor’ or V ‘bad’, indicating that invertebrate communities were negatively affected by pesticide inputs. Regarding stream hydromorphology, monitoring results revealed that 65% of the sampled agricultural stream sites did not achieve good ecological status. Citizen science data for macroinvertebrate communities (bioindicator SPEAR_pesticides_) and stream hydromorphology showed a high level of agreement with professional assessments.

These validated citizen science data on small streams are suitable to complement official WFD reporting by environmental agencies on the ecological status of freshwater ecosystems. By involving diverse stakeholders such as environmental NGO groups, anglers and high schools in tracking land use impacts and restoration efforts in small streams, citizen science stream monitoring can support local decision making for sustainable water management and stream protection.

### 4.1 Accuracy of citizen science stream monitoring data

We have demonstrated that, with appropriate training and guidance, citizen scientists are able to provide valid data on stream hydromorphology and on benthic invertebrate community composition as an indicator for pesticide exposure applying the bioindicator SPEAR_pesticides_. As identified in a previous study, data accuracy in citizen science stream monitoring depends on the taxonomic level analyzed and the commonness of the different invertebrate taxa (von Gönner et al., 2023a). The FLOW methodology works primarily at the family level, with a small percentage of specimens identified at the genus level by experienced citizen scientists, or at the order level, e.g. if specimens are damaged. The proportion of correctly identified invertebrates was high at family level in the majority of CS samples analyzed in this study (84% on average), and we found a highly significant relationship (R^2^ = 0.79) between CS and professional SPEAR_pesticides_ values. However, in 17% of the CS samples, the proportion of correct CS family identifications was relatively low (<70%, see Fig.1), likely due to lower levels of experience or motivation among the respective CS groups. Nevertheless, the overall high agreement between CS and professional SPEAR_pesticides_ values indicates that the SPEAR index can provide a useful assessment of invertebrate community composition, even when based on coarser functional trait data (comparable to order-level data, Liebmann et al., 2022). SPEAR_pesticides_ values provided by citizen scientists averaged slightly higher than professional SPEAR values because some common invertebrate taxa are classified as pesticide-sensitive at the family level (e.g., Limnephilidae, Goeridae) but are classified as pesticide-insensitive at the genus or species levels (e.g. Anabolia sp., Silo sp.), which were mostly not recorded by citizen scientists. All in all, this study demonstrates that trained citizen scientists can provide valid information on macroinvertebrate community composition in different ecological conditions (see also von Gönner et al., 2023a; Storey et al., 2016).

The citizen scientists’ hydromorphology status class assessments agreed with those of the professional ecologists in two thirds of the sample sites. This exceeds the results of our previous study with a smaller sample of n = 28 stream sites (von Gönner et al., 2023a). Although the citizen science and professional site classifications into good or unsatisfactory hydromorphological status were consistent for 85% of the sample sites in the present study, we observed that citizen scientists tended to rate stream hydromorphology more often as ‘moderate’ then professional ecologists did (Fig. 1B), which we still estimate to be in acceptable observer error margins. We acknowledge that it may take some field experience and practice at different stream sites to arrive at more consistent ecological assessments, which citizen scientists may acquire over time.

Citizen science point measurements of physico-chemical water parameters should only be considered as a snapshot of water quality on the respective citizen science stream sampling days, mainly due to the limited measurement frequency (see section 4.2. below, von Gönner et al., 2023a).

Overall, the generation of scientifically valid citizen science freshwater monitoring data requires hands-on training and feedback for citizen scientists and, in the case of novices, assistance with fieldwork by experienced biologists (von Gönner et al., 2023a). As demonstrated by the evaluation of citizen science data accuracy in this study, multiplier training is an appropriate method to adequately prepare a large number of spatially dispersed citizen science groups for standardized stream monitoring.

### 4.2 Ecological status of streams

Our sample of 137 stream sample sites covers a gradient of agricultural land use and a wide range of stream types in different ecoregions. Therefore, the present citizen science monitoring results provide comprehensive insights into the status of small streams in Germany and are a valuable complement to the WDF monitoring, which focuses on large rivers and streams.

The citizen science monitoring data from our study corroborate the results of previous studies on the poor ecological status of rivers (e.g. EEA, 2018; UBA, 2022) and small water bodies (Liess et al., 2021).

The SPEAR_pesticides_ bioindicator indicates that macroinvertebrate communities in 58% of the agricultural streams investigated are adversely affected by pesticide residues (corresponding to a moderate to bad SPEAR_pesticides_ class). This is in line with the results of a recent survey of small water bodies representative for the gradient of agricultural land use in Germany, which found that 83% of agricultural streams did not meet ecological targets related to pesticides and 81% of sites had pesticide concentrations exceeding the regulatory acceptable concentrations (Liess et al., 2021). The lower proportion of streams severely affected by pesticides in our study may be attributed to a relatively lower proportion of agricultural land cover in the agricultural stream site catchments investigated here (mean = 62% compared to 77% in Liess et al., 2021). Similar to previous studies (Liess et al., 2021; Szöcs et al., 2017), our results show that pesticide pressure (here quantified by the SPEAR_pesticides_ bioindicator) increased with the share of agricultural land use in the catchments of the sampled streams sites (Fig. 4).

We also found that stream hydromorphology was strongly affected by human activities in 65% of the agricultural and in 58% of the non-agricultural stream sites, in line with previous monitoring results of poor habitat quality in many Central European rivers and streams (Lorenz et al., 2004, 2009; UBA, 2022). In particular, substrate diversity, riparian vegetation, and longitudinal profile including flow diversity and habitat continuity, which are essential for many benthic invertebrates and (semi)-aquatic vertebrates, were often classified as severely altered or depleted by the citizen scientists (SI Fig.3).

The distributions of physico-chemical measurements show, similar to previous monitoring results on rivers (Poikane et al., 2021; Sadayappan et al., 2022; UBA, 2023), that a majority of the sampled streams are affected by high nutrient concentrations (especially phosphate and nitrate). This eutrophication can promote excessive algal growth, which can lead to oxygen depletion and harm aquatic organisms (Hilton et al., 2006). However, the citizen science point measurements of physico-chemical parameters should be interpreted as supplemental information only, as single measurements are of limited significance and may under- or overestimate exceedance frequencies of annual thresholds. For a representative analysis of physico-chemical status, the frequency of citizen science measurements should be increased and citizen science test kits should be regularly calibrated with field methods and environmental agency test procedures (Quinlivan et al., 2020; von Gönner et al., 2023a).

Another limitation of our study is that the citizen science project FLOW does not investigate all endpoints for assessing overall ecological status under the WFD (macrophytes, algae and fish are not included), and the bioindicator SPEAR_pesticides_ assesses invertebrate communities specifically in relation to pesticide exposure. Therefore, our results are not directly comparable with the results of the German Environment Agency (UBA, 2022), which shows that only 8% of the official stream sampling sites are in good condition.

Numerous insects depend on ecologically intact streams with good water and habitat quality (Dijkstra et al., 2014). For example, many sensitive mayfly, stonefly and caddisfly larvae only tolerate low levels of toxic pesticides or oxygen depletion and require a structurally rich streambed with a variety of substrates such as dead wood and gravel of varying sizes for feeding and reproduction. Streams in poor ecological condition therefore cannot fulfill their function as ‘lifelines’ in the intensively used cultural landscape and their contribution to the preservation of biodiversity.

The poor ecological status of surface waters severely compromises ecosystem services that are essential for human well-being: high quality water supply and storage, filtering function and flood protection provided by near-natural floodplains, conservation of freshwater biodiversity, as well as the provision of cultural services such as recreation in near- or semi-natural, aesthetic landscapes (Böck et al., 2018). This affects all sectors of society from private households to companies, municipalities and federal states (BMUV, 2023).

Even more than 20 years after the adoption of the WFD, the EU member states are still far from implementing the WFD’s goal to conserve and restore the good ecological status of surface waters. This has been attributed to difficulties with reconciling multiple conflicting user interests and integrating different water-related policies (e.g., water supply, flood control, agriculture, recreation, nature protection); a lack of funding and land availability for stream restoration; unclear responsibilities for implementing river basin management plans; and too little cooperation between environmental agencies, water associations, NGOs, and citizen initiatives (Reese et al., 2018; Carvalho et al., 2019). In addition, the current WFD monitoring design has been shown to be unsuitable for detecting pesticide risks in small streams (Weisner et al., 2022), as it is focused on larger water bodies and limited by a too small range of substances investigated. This deficit in monitoring becomes particularly critical in light of the European Green Deal’s goal to halve the amount and risk of pesticide use by 2030. To address this issue, citizen science could be an effective means to expand the spatial and temporal scale of WFD freshwater monitoring. Targeted fine-grain citizen science monitoring could be used as a ‘screening monitoring’ to identify water bodies strongly affected by pesticide inputs or morphological degradation. Based on citizen science results, more in-depth monitoring could be conducted and appropriate mitigation measures (see section 4.3 below) could be designed and implemented together with local citizen groups.

### 4.3 Conclusions and implications for policy and practice

Our study highlights the desolate ecological status of small streams in German agricultural landscapes, affected by poor hydromorphology and severely impoverished benthic invertebrate communities, indicating high pesticide pressures. We could show that pesticide-induced adverse effects on benthic invertebrate communities, as quantified by the SPEAR_pesiticides_ indicator, correlate with the proportion of agricultural land in the stream catchments.

The present results of the citizen science stream monitoring program FLOW underscore the urgency of making progress in freshwater protection. They highlight the need for rapid implementation of the Water Framework Directive, the Nature Restoration Law and the Sustainable Use Regulation at EU level to protect aquatic biodiversity and freshwater ecosystems from pesticide exposure and habitat degradation.

Our monitoring results also emphasize the need to generate societal, medial and political attention for freshwater protection. By integrating citizen expertise into freshwater research and by offering opportunities for networking and community-building, citizen science projects can encourage local ownership for freshwater protection and restoration (Brooks et al., 2019). Citizen science freshwater monitoring could thus become an important tool to raise public awareness, foster collective efficacy (von Gönner et al. 2023b) and initiate community-based stewardship for stream health (Huddart et al., 2016). In this way, engaged citizens could help ensure that national and international freshwater conservation targets are increasingly recognized and implemented at the local and regional level.

In several cases, engaged citizens have already helped identify and report water pollution to authorities, resulting in the implementation of water protection measures (e.g. Flint water crisis, Pieper et al. 2018; UK Angler’s Riverfly Monitoring Initiative, Brooks et al. 2019). Likewise, decision makers and media professionals could use the citizen science-driven evidence of poor small stream health presented here to strategically advance freshwater protection with concrete action programs.

Experience with the WFD shows that, in addition to extensive monitoring efforts, locally adapted, participatory planning and implementation processes are needed to restore the good ecological status of rivers and streams (Edwards et al., 2018; Carvalho et al., 2019; European Commission 2019b). Combined with ongoing citizen science monitoring, the implementation of stream restoration measures should also involve citizen initiatives and various relevant stakeholders, thereby strengthening the social license for conservation (Kelly et al., 2019). In Germany, numerous citizen groups engaged in the FLOW project are motivated to take action in the field of stream restoration based on their citizen science monitoring results. The poor ecological status of many streams could be improved with low-threshold measures, which can also be initiated by local citizen initiatives, as they generally do not require an extensive planning approval procedure (UBA, 2019). These include, for example, the planting of native, site-specific shrubs and trees in the riparian zone to provide shade and create a buffer against runoff from agricultural fields (vegetated buffer strips, see Vormeier et al. 2023); or the small-scale introduction of gravel or dead wood to increase substrate and flow diversity in streams (Madsen and Tent, 2000).

To enhance the scientific and political impact of citizen science freshwater monitoring in tandem with official monitoring, future research is needed to identify targeted criteria for citizen science sample site selection. In this way, citizen science monitoring could specifically fill gaps in official monitoring programs and monitor catchments with a high risk of pesticide exposure or morphological degradation, or catchments for which no official monitoring data exist. Citizen science freshwater monitoring initiatives should be actively supported by research policy and societal decision makers to enable and foster collective freshwater stewardship. Given the state of freshwater streams with the majority of small agricultural streams in poor ecological status, effective restoration and appropriate mitigation measures are needed to meet national and international targets for functioning freshwater ecosystems. In this way, Germany and Europe could take a major step forward in maintaining and restoring the health of rivers and streams.

## Acknowledgements

This study could only be realized thanks to the engagement and expertise of over 900 citizen scientists across Germany who joined the FLOW project to explore their local streams. Special thanks to Anna-Katharina Klauer and the Saxonian Foundation for Nature and Environment (LaNU) for cooperating with their mobile lab ‘Umweltmobil Planaria’. The data collection would not have been possible without the development of the FLOW web application coordinated by Alexander Harpke, Daniel Sielaff and Jan Bumberger from the Research Data Management Team of Helmholtz Centre for Environmental Research – UFZ in cooperation with InfAI (Institut für Angewandte Informatik). The FLOW project was funded by the German Federal Ministry of Education and Research (BMBF, Project Grant 01BF1906). Additional support was provided by a PhD scholarship from the Deutsche Bundesstiftung Umwelt (DBU, AZ 20019/592) and by the project “Belastung von kleinen Gewässern in der Agrarlandschaft mit Pflanzenschutzmittel-Rückständen - TV1 Datenanalyse zur Pilotstudie Kleingewässermonitoring 2018/2019” funded by the German Federal Ministry for the Environment, Nature Conservation, Nuclear Safety and Consumer Protection (BMUV, FKZ 3720 67 4011). We also acknowledge support from iDiv funded by the German Research Foundation (DFG-FZT 118, 202548816).

## Author Contributions

**Julia von Gönner:** Conceptualization, Methodology, Validation, Formal analysis, Investigation, Writing – Original Draft, Writing – Review & Editing; **Jonas Gröning:** Conceptualization, Data curation, Validation, Formal analysis, Visualization, Writing – Original Draft, Writing – Review & Editing; **Lilian Neuer:** Conceptualization, Investigation, Resources; **Volker Grescho:** Software, Data curation; **Veit Hänsch, Benjamin Laue, Eva Molsberger-Lange, Elke Wilharm:** Investigation, Writing – Review & Editing; **Matthias Liess:** Supervision, Conceptualization, Writing – Review & Editing; **Aletta Bonn:** Supervision, Conceptualization, Writing – Review & Editing.

## Data availability

The data analyzed in this study is available upon request and will be archived on Pangaea (title: “Stream monitoring data from the citizen science project FLOW, 2021-2023”, embargo until publication).

