## Supplementary Information for "The majority of Germany’s small agricultural streams are in poor ecological status"

**SI Fig. 1: Distribution of A. catchment areas, B. agricultural area, C. arable land and D. urban land cover in catchments of citizen science stream sample sites.**

A: Histogram of catchment area size of the citizen science stream sample sites (n=137). Three sites with catchment areas > 100km$^2$ are not displayed.

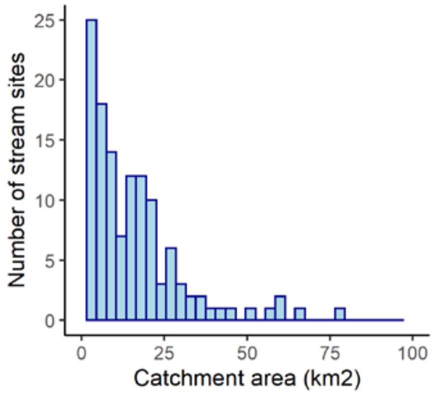


B: Histogram of percentage of agricultural land cover of the citizen science stream sample sites (n=137).

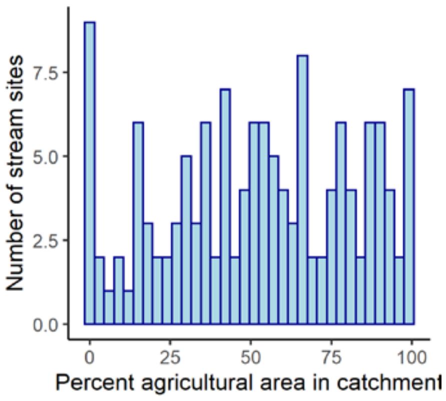


C: Histogram of percentage of arable land cover of the citizen science stream sample sites (n=137).

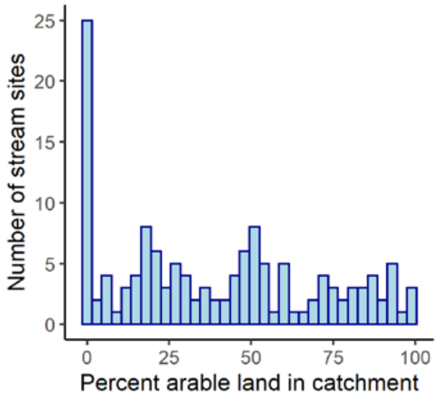


D: Histogram of percentage of urban land cover of the citizen science stream sample sites (n=137).

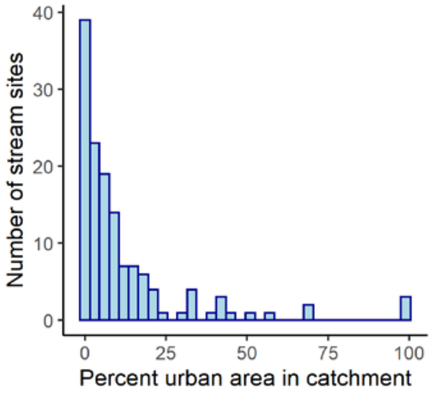

**SI Tab. 1: Distribution of stream types among the n=137 sample sites analyzed.** Stream types are listed according to German river typology, Pottgiesser 2018; LAWA 2019. For details on sampling dates and site-specific information, see data supplement on Pangaea (embargo until publication of manuscript).

| Stream type | Number of stream sites | Percent of stream sites |
|---|---|---|
| Small coarse substrate dominated siliceous highland rivers (type 5) | 28 | 20.4% |
| Small fine substrate dominated calcareous highland rivers (type 6) | 22 | 16.1% |
| Small sand-dominated lowland rivers (type 14) | 19 | 13.9% |
| Small loess and loam-dominated lowland rivers (type 18) | 15 | 10.9% |
| Small coarse substrate dominated calcareous highland rivers (type 7) | 10 | 7.3% |
| Small organic substrate-dominated rivers (type 11) | 10 | 7.3% |
| Small gravel-dominated lowland rivers (type 16) | 9 | 6.6% |
| Mid-sized fine to coarse substrate dominated siliceous highland rivers (type 9) | 6 | 4.4% |
| Streams in the alpine foothills (type 2) | 5 | 3.6% |
| Small fine substrate dominated siliceous highland rivers (type 5.1) | 4 | 2.9% |
| Small streams in riverine floodplains (type 19) | 4 | 2.9% |
| Mid-sized fine to coarse substrate dominated calcareous highland rivers (type 9.1) | 2 | 1.5% |
| Mid-sized and large sand and loam-dominated lowland rivers (type 15) | 2 | 1.5% |
| Streams in the Pleistocene sediments of the alpine foothills (type 3) | 1 | 0.7% |

**SI Tab. 2: Overview of citizen science training material used in this study (in German language).**

| No. | Material | Access (can be viewed at this URL): |
|---|---|---|
| 1 | FLOW project booklet (65 pages) | |
| 2 | FLOW macroinvertebrate identification booklet (84 pages) | |
| 3 | Video tutorials on FLOW monitoring methods (in German).<br>Note: As macroinvertebrate sampling requires an official permission from the state environmental agencies in Germany, the videos are only intended for registered project participants, please do not distribute. | |
| 4 | Online quiz to practice macroinvertebrate identification and assessment of stream hydromorphology | |

### SI Tab. 3: Overview of citizen science physico-chemical measurement methods.

| **Physicochemical water parameter** | **Measuring method and equipment** |
|---|---|
| Dissolved oxygen (mg/l) | Colorimetric oxygen test kit Visocolor Eco, Macherey-Nagel, Düren, Germany |
| Water temperature (°C)<br>Electrical conductivity (µS/cm) | Thermometer and conductivity meter: HM Digital Inc., TDS EC-3 |
| Flow velocity (m/sec) | “Floating body” method: the distance (m) that a floating object (stick) passed within 10 seconds was measured with a measuring tape, flow velocity was calculated as meters per second |
| Ammonium (mg/l $NH_4$)<br>Phosphate (mg/l $PO_4$)<br>Nitrite (mg/l $NO_2$-)<br>Nitrate (mg/l $NO_3$-)<br>pH | Colorimetric tests MQuant/ MColortest, Merck KGaA, Darmstadt, Germany<br>The water sample was prepared according to instructions with indicator reagents. Then, the color of the water sample was compared to the respective colorimetric color chart, which provided a specific measuring value for each color point. If the color of the water sample fell between two color points, the two corresponding measuring values were averaged. |

**SI Tab. 4: Summary of citizen science SPEAR$_{pesticides}$ and hydromorphology index values in 2021, 2022 and 2023.** For the analysis of SPEAR$_{pesticides}$, we excluded all stream sites monitored by the citizens in the respective year that showed a very low flow velocity (< 0.05 m/sec, see Liess et al. 2021).

| Year | Mean (sd) SPEAR index | Year | Mean (sd) Hydromorphology index |
|---|---|---|---|
| 2021 (n=34) | 0.44 (0.23) | 2021 (n=34) | 4.04 (0.98) |
| 2022 (n=52) | 0.63 (0.35) | 2022 (n=62) | 3.71 (1.10) |
| 2023 (n=65) | 0.63 (0.28) | 2023 (n=76) | 3.82 (1.11) |

**SI Tab. 5: Percentages of SPEAR$_{pesticides}$ and hydromorphology status classes recorded by the citizen scientists in the three monitoring years 2021, 2022 and 2023.** This summary includes all stream sites monitored by citizen scientists in the given year, including data points for stream sites that were also monitored in the previous or subsequent year (n=172, the total sample analyzed in the manuscript of n=137 sites includes only the most recent data point per stream site). For the analysis of SPEAR$_{pesticides}$ in this summary, we excluded all stream sites monitored by the citizens in the respective year that showed a very low flow velocity (< 0.5 m/sec).

| SPEAR$_{pesticides}$ class | 2021 (n=34) | 2022 (n=52) | 2023 (n= 65) |
|---|---|---|---|
| I – high | 2.9% | 32.7% | 26.2% |
| II – good | 23.5% | 15.4% | 26.2% |
| III – moderate | 35.3% | 21.2% | 23.1% |
| IV – poor | 20.6% | 21.2% | 18.5% |
| V – bad | 17.6% | 9.6% | 6.2% |

| Hydromorphology class | 2021 (n=34) | 2022 (n=62) | 2023 (n=76) |
|---|---|---|---|
| I – unchanged | 2.9% | 4.8% | 7.9% |
| II – slightly changed | 32.4% | 35.5% | 30.3% |
| III – moderately changed | 35.3% | 37.1% | 32.9% |
| IV – strongly changed | 23.5% | 19.4% | 23.7% |
| V – completely changed | 5.9% | 3.2% | 5.3% |

**SI Fig. 2: Maps of Germany showing citizen science SPEAR$_{pesticides}$ and hydromorphology index values recorded in the three monitoring years 2021, 2022 and 2023.** For SPEAR$_{pesticides}$ we excluded all stream sites monitored by the citizens in the respective year that showed a very low flow velocity (< 0.05 m/sec).

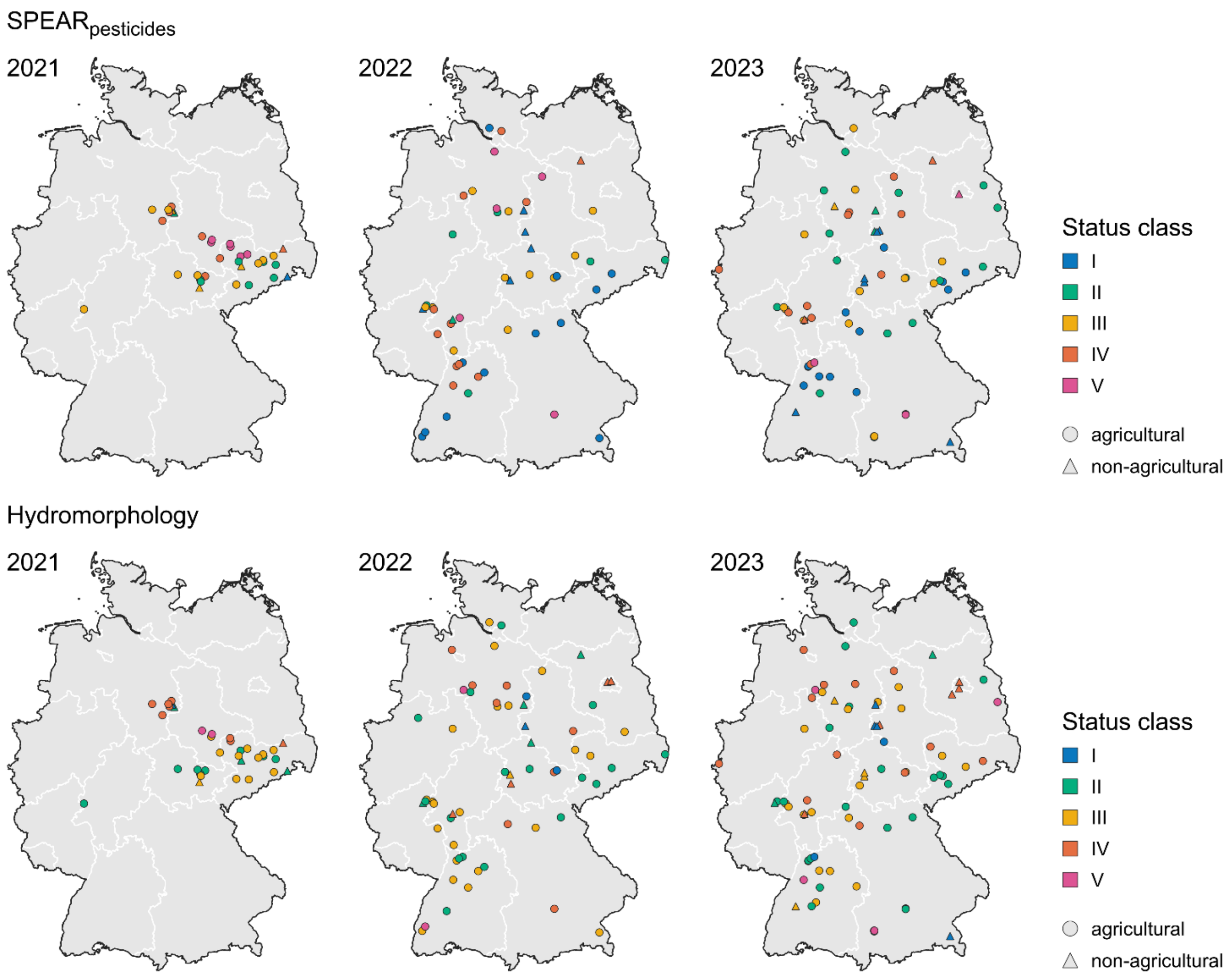

**SI Fig. 3: Overview of citizen science status classifications of the six hydromorphology subcomponents at n=113 agricultural and n=24 non-agricultural stream sites.**

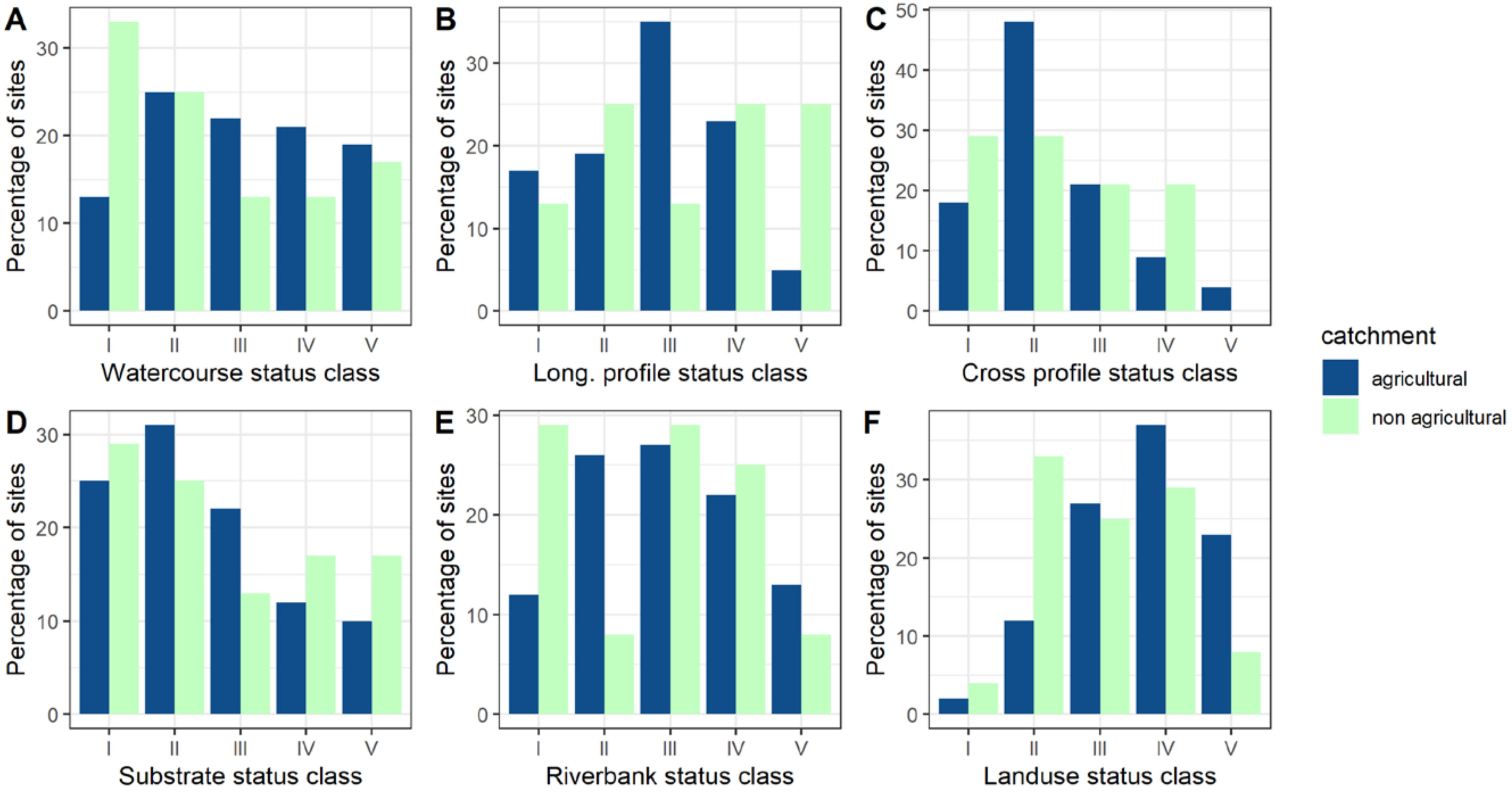

**SI Fig. 4: Citizen science and professional classification of stream sites into (A) $SPEAR_{pesticides}$ status classes and (B) ecological status according to $SPEAR_{pesticides}$ (good or unsatisfactory on a scale of 0 to 1; n = 81 sites).** 64% of the sites were classified into the same status class by citizen scientists and professionals (Fig. 4.A), and both monitoring teams reached the same conclusion about ecological status in 84% of the cases (Fig. 4.B).

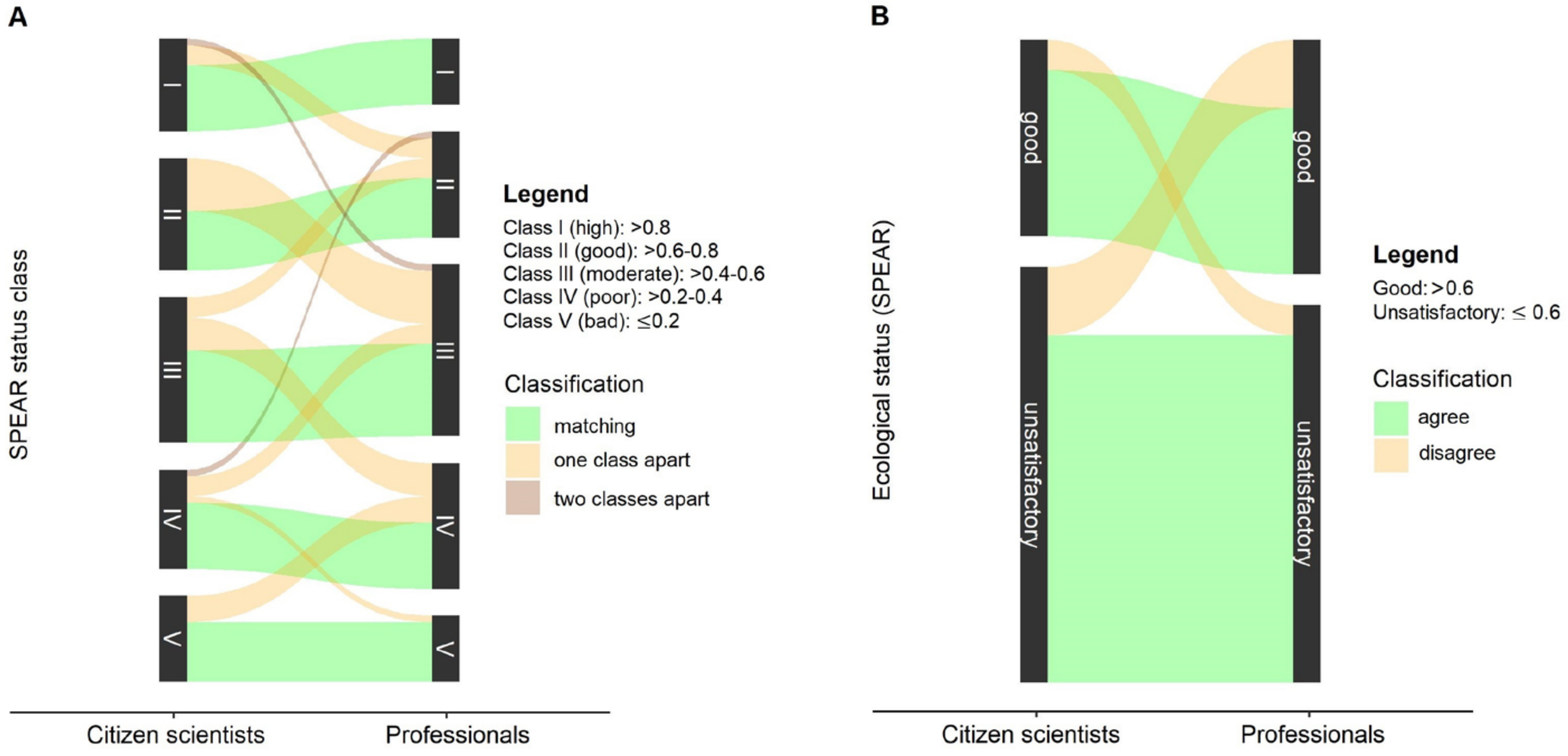

**SI Fig. 5: Citizen science and professional classification of stream sites into (A) hydromorphology status classes and (B) hydromorphological status (good or unsatisfactory; n = 79 sites).** 65% of the sites were classified into the same status class by citizen scientists and professionals (Fig. 5.A), and both monitoring teams reached the same conclusion about ecological status in 85% of the cases (Fig. 5.B).

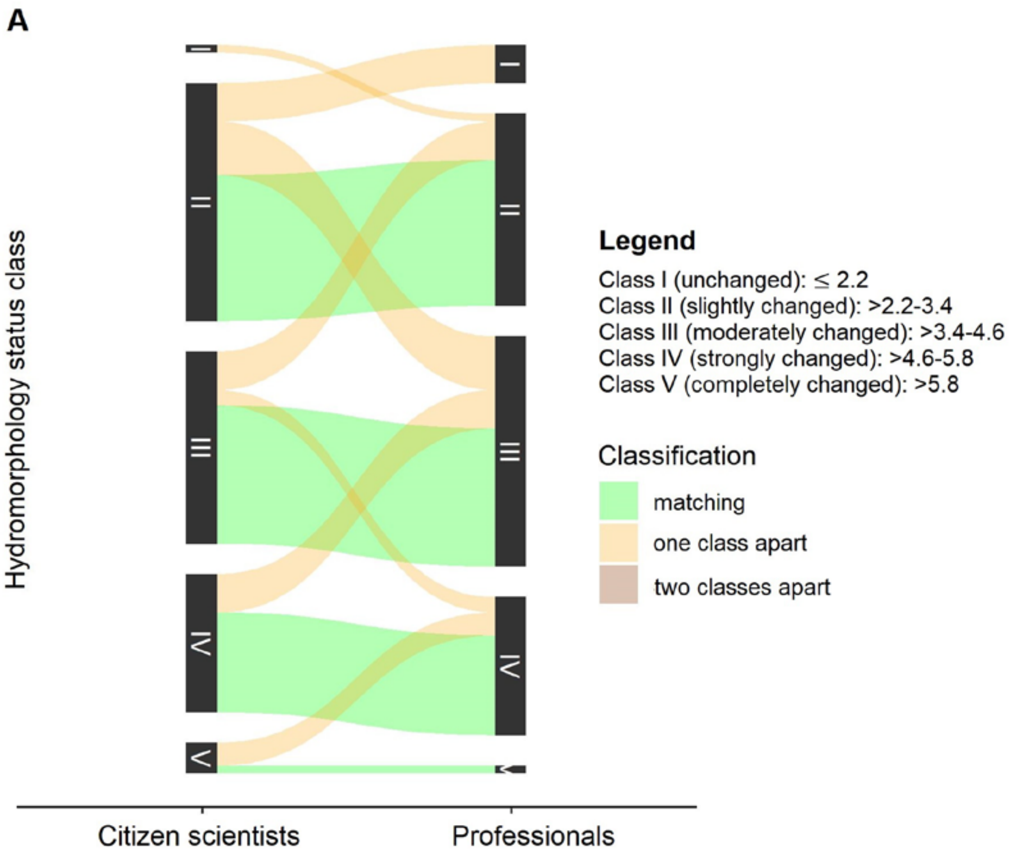


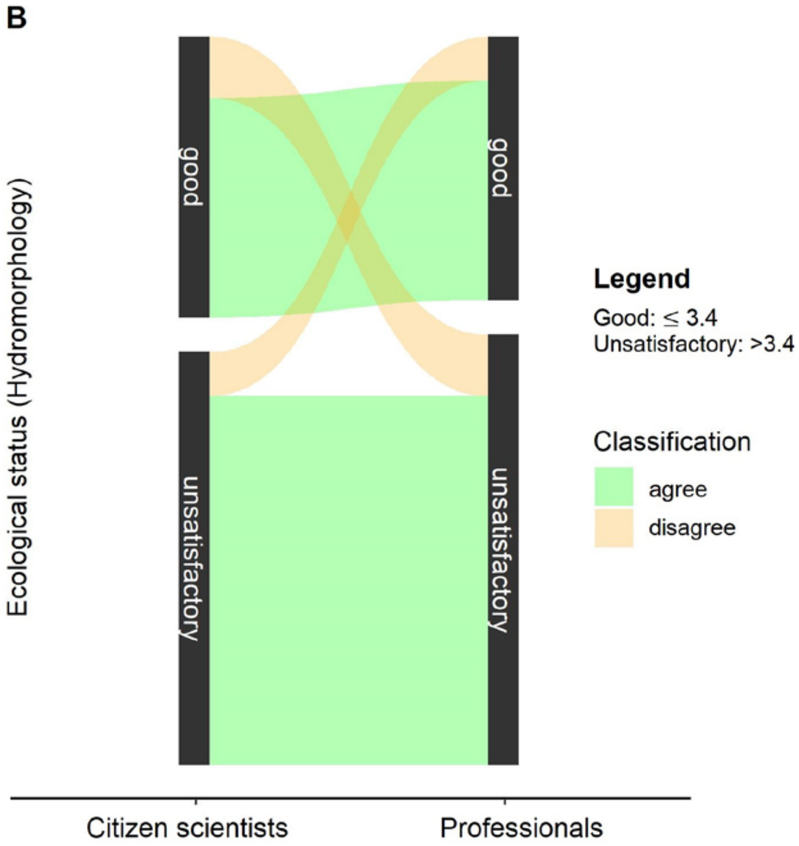
